# Stress fiber growth and remodeling determines cellular morphomechanics under uniaxial cyclic stretch

**DOI:** 10.1101/622092

**Authors:** Aritra Chatterjee, Paturu Kondaiah, Namrata Gundiah

## Abstract

Stress fibers in the cytoskeleton are essential in maintaining cellular shape, and influence their adhesion and migration. Cyclic uniaxial stretching results in cellular reorientation orthogonal to the applied stretch direction *via* a strain avoidance reaction; the mechanistic cues in cellular mechanosensitivity to this response are currently underexplored. We show stretch induced stress fiber lengthening, their realignment and increased cortical actin in fibroblasts stretched over varied amplitudes and durations. Higher amounts of actin and alignment of stress fibers were accompanied with an increase in the effective elastic modulus of cells. Microtubules did not contribute to the measured stiffness or reorientation response but were essential to the nuclear reorientation. We modeled stress fiber growth and reorientation dynamics using a nonlinear, orthotropic, fiber-reinforced continuum representation of the cell. The model predicts the observed fibroblast morphology and increased cellular stiffness under uniaxial cyclic stretch. These studies are important in exploring the differences underlying mechanotransduction and cellular contractility under stretch.

## Introduction

Adherent fibroblasts in tissues, such as arteries and lungs, undergo cyclic stretch and respond to changes in their mechanical milieu through a complex interplay between the cytoskeletal components and integrin-mediated pathways to influence cell contractility and secretion of extracellular matrix constituents [1,2]. Cell driven biochemical processes result in gain or loss in mass and induce remodeling of the underlying material properties such as stiffness and anisotropy. Continuum mechanics-based approaches to address biological growth and remodeling suggest an intimate relationship between these processes which results in non-uniform changes to overall form and function of the underlying structures [3]. How do mechanosensing processes influence cellular growth and remodeling under stretch? Stretch induced reorientation of cells involves both passive mechanical response to cyclic substrate deformation and dynamic changes to the cytoskeleton [4]. Contractile stress fibers (SF), comprised primarily of actin and myosin II, are essential in the development of intracellular stresses which are transmitted through focal adhesion (FA) complexes and depend on the activity of RhoA, active Rac1, or Cdc42 [5–7].

Cyclic uniaxial stretching of fibroblasts seeded on isotropic elastomeric substrates leads to cell reorientation from random to a near perpendicular angle to the loading direction [8,9]. Vascular smooth muscle cells under stretch show an initial rapid growth and reinforcement followed by disassembly of SF aligned in the direction of stretch; the SF subsequently rearrange in an orthogonal direction [10]. The associated actin reorientation under uniaxial stretch is accompanied with an increase in FAK signaling, MAP kinases, and a corresponding dynamic increase in the size of FA complexes [11]. Recent experiments show reorganization and remodeling of FA due to stretching of cells [11–13]. Higher strain rates caused rapid disassembly of SF’s along the direction of deformation [14,15]. The application of cyclic uniaxial strain also influence cell spreading on soft substrates; SF formation persisted four hours after stretching and was related to temporal changes in the movement of MRTF-A (myocardin-related transcription factor-A) and YAP (Yes-associated protein) from the cytoplasm to the nucleus [13,16].

Stress fibers are exquisitely sensitive to the dynamically changing mechanical environment and are essential in the regulation of cell polarization and transmission of tensional force from the FA to the cell [17]. Cell spreading, dependent on the stiffness of the underlying substrate, resulted in a bi-phasic cellular orientation caused by *de novo* formation of ventrally oriented actin in the cell. Cell alignment under cyclic uniaxial stretch is hypothesized to be an avoidance reaction to stretch which is facilitated *via* cell-substrate interactions and their links to the cytoskeleton [18]. The mean cell orientation after stretch contributes to the tensional homeostasis of the cell and helps maintain an optimal internal stress [19]. More recently, Chen *et al* showed that traction boundary conditions are crucial in the determination of cellular orientation under stretch [20] The precise mechanistic cues in cellular mechanosensitivity to dynamical environments and links to stress fiber growth and cell stiffness changes however remains a fundamental open question.

The goals of this study are to investigate SF dynamics in fibroblasts under uniaxial cyclic stretch and relate their growth and remodeling to the cellular reorientation response using a morphoelasticity framework. We used a custom fabricated stretcher to subject fibroblasts to uniaxial cyclic stretch for different amplitudes over varied time periods. We show that the lengthening and realignment of SF primarily contributes to the cellular reorientation response under cyclic stretch. We model these morphomechanical responses using nonlinear, hyperelastic, fiber-reinforced material model for the cell comprising of two families of SF that grow and reorient to under uniaxial cyclic stretch. The measured changes in cell stiffness in response to the stretch amplitude and time duration due to SF reorientation are well captured using the model. These results show the importance of SF growth in the resulting elongation and orientation response of cells under cyclic uniaxial stretch that are important in the overall mechanotransduction and cellular contractility.

## Results

### Cyclic stretch induced changes in cell morphology and stiffness depends on SF reorientation and cortical actin thickness

Uniaxial cyclic stretch experiments were performed on NIH 3T3 fibroblasts seeded on thin flexible membranes (20 mm length x 10 mm width x ~150 µm thickness), fabricated from poly dimethyl siloxane (PDMS; Sylgard®184, Dow Corning) mixed in the ratio of 10:1, using a custom stretcher (Supplementary Fig. S1). Cell seeded constructs were stretched uniaxially at 5% and 10% amplitude at a frequency of 1 Hz for 3 and 6 hours. Confocal images of fibroblasts stretched at 10% amplitude for 6 hours (A10T6) show distinct changes in the overall cellular morphology. Cells were significantly elongated and elliptical along a uniform direction perpendicular to the direction of applied stretch as compared to unstretched controls (Fig. 1). There were clear changes that were apparent in the cytoskeletal and nuclear orientations at higher amplitude and over longer stretching durations (A10T6) (Fig. 2a, b). These results show significant differences in the major axis length of the cell along the direction of reorientation and in cellular aspect ratios (p<0.01 for both) between cyclically stretched cells (A10T6) and corresponding unstretched controls (Fig. 2d, e). In addition, the cell spread area increased significantly after subjecting to cyclic stretch (3087±854 µm^2^) as compared to unstretched controls (2450 ±357 µm^2^; p<0.05). We quantified the SF orientation distributions, based on a wedge-shaped orientation filter in MATLAB (MathWorks 2016b), from immunofluorescence images of cells (n=30) stained to visualize SF (Supplementary Fig. S2) [21,22]. SF lengths were quantified using Hough transform. A total of ~30 cells were included in each set and the data averaged to compute an overall SF angular and length distribution (Table 1). The results show SF realignment in a direction near perpendicular to the direction of applied cyclic stretch; nuclear orientation was also in the direction of cytoskeletal reorientation (Fig. 2c). SF orientations changed from a bimodal distribution in control unstretched cells to a unimodal distribution orthogonal to the stretch direction due to uniaxial cyclic stretch. The SF distribution showed a distinct peak at higher amplitude and stretch duration (A10T6). These data suggest that almost all cells oriented to a uniform near perpendicular angle to the direction of applied cyclic stretch.

**Figure 1.**
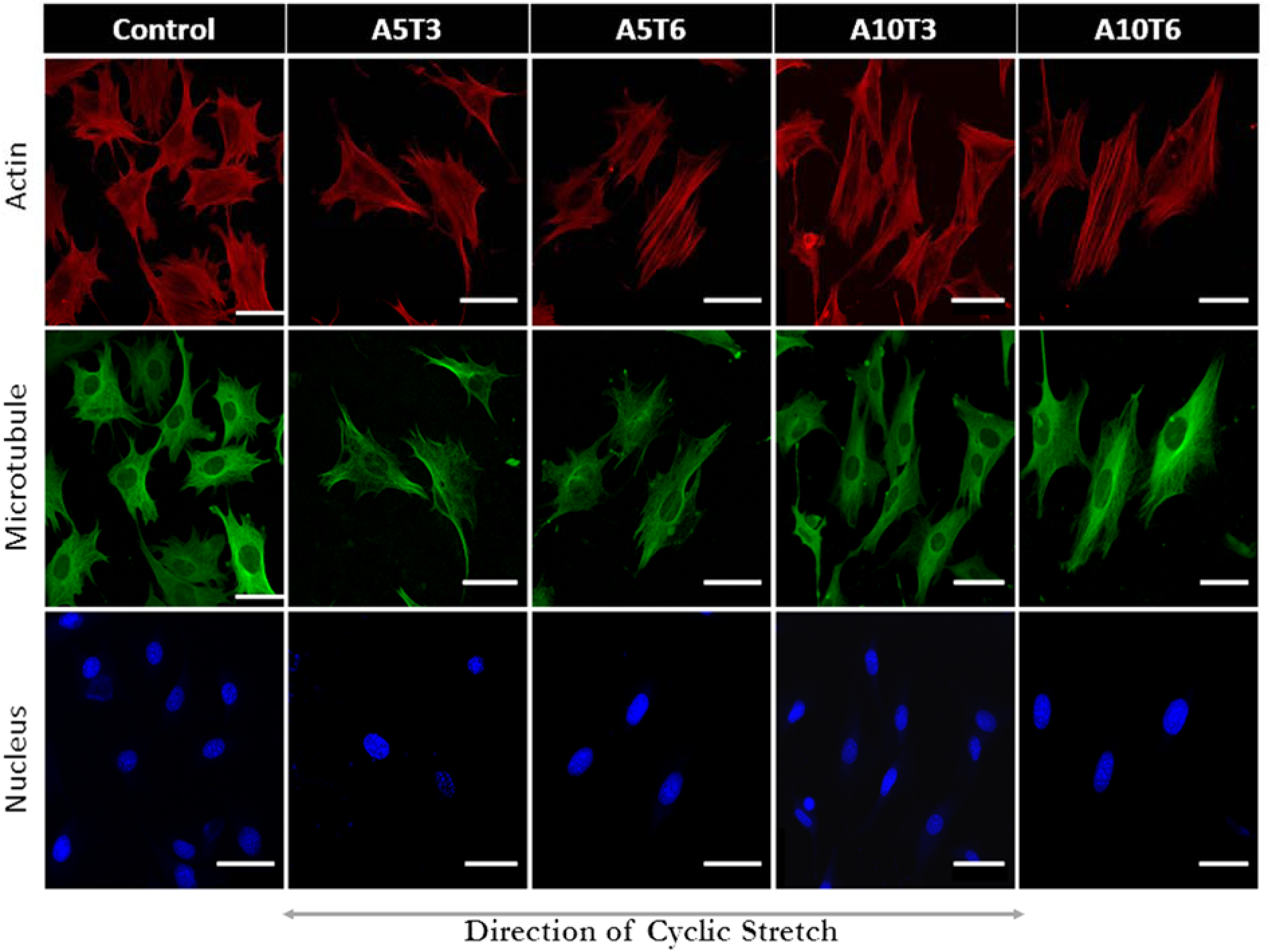
Fibroblast cells, cultured on PDMS membranes and subjected to uniaxial cyclic stretch (1 Hz) for different durations, were fixed and imaged to visualize actin (red), microtubules staining (green), and the nuclei (blue). The direction of cyclic uniaxial stretch is indicated using an arrow. Stretch amplitude (X % elongation) and duration of the experiment (Y in hrs) are represented as AXTY. Unstretched cells on PDMC membranes were used as controls in the study. Scale bar represents 50 μm.

**Figure 2.**
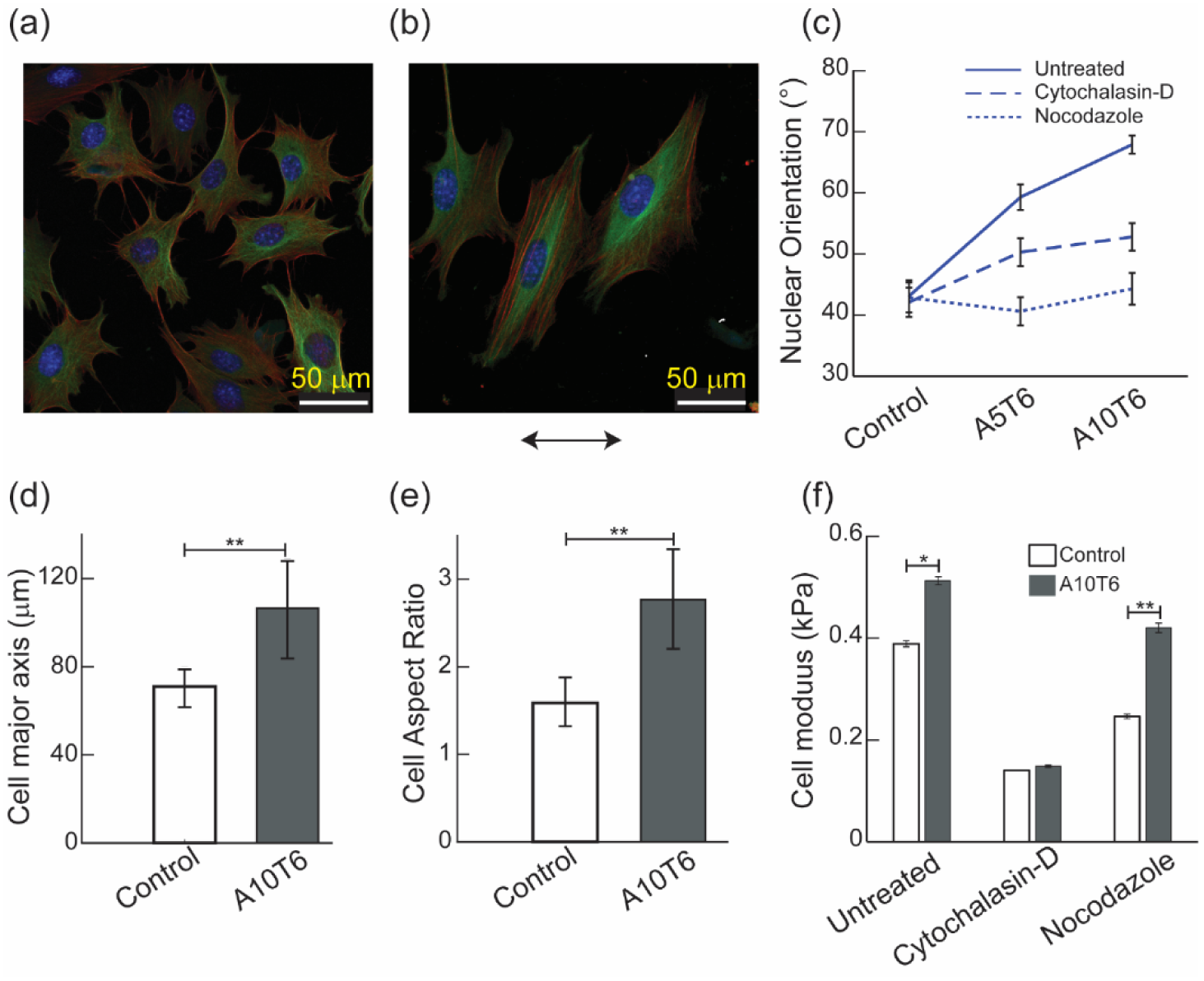
**a.** F-actin (red), microtubule (green) and DAPI nucleus (blue) stained merged confocal images of control fibroblasts on unstretched PDMS membranes. **b.** Fibroblasts on unidirectionally stretched PDMS membranes at 10% amplitude for 6 hours (A10T6). **c.** The angular orientation of the nucleus was quantified under unidirectional cyclic stretch (1 Hz) with/ without cytochalasin-D and nocodazole. No significant nuclear reorientation was observed with depolymerization of microtubules; the re-orientational response was also reduced in cells following actin depolymerization **d**. Cyclic stretch (A10T6) induced fibroblast elongation was quantified using the major axis of the cell. **e**. The cell aspect ratio increased under unidirectional cyclic stretch. **f**. AFM was used to measure the effective elastic modulus of cells, subjected to cyclic stretch in the presence and absence of cytoskeletal disruptors, and are shown for control cells (unstretched and un-treated) and cells stretched using 10% amplitude for 6 hours (A10T6; n=25 each group). Significant differences between the different groups are indicated for p<0.01 (**) and p<0.05 (*).

**Table I:** Distribution of the angular orientations (Mean ± SEM) and lengths (Mean ± SEM) are shown for fibroblasts on PDMS membranes and subjected to uniaxial cyclic stretch (1 Hz). The stretch amplitude (X % elongation) and duration of the experiment (Y in hrs) are represented as AXTY.

|  | ***SF angles (°)*** | ***SF length (μm)*** |
| --- | --- | --- |
|  | ***(Mean ± SEM)*** | ***(Mean ± SEM)*** |
| Control | 43.6 ± 4.9 & 150.9 ± 7.9 | 25.1 ± 1.6 |
| A5T3 | 51.8 ± 5.3 & 148.9 ± 4.9 | 28.3 ± 1.7 |
| A5T6 | 76.2 ± 7.5 & 127.5 ± 5.5 | 32.5 ± 2.2 |
| A10T3 | 72.7 ± 7.6 & 115.2 ± 7.3 | 31.1 ± 1.7 |
| A10T6 | 79.9 ± 2.8 & 99.2 ± 1.4 | 58.5 ± 3.6 |

We used cytoskeletal inhibitors to delineate the individual contributions of SF’s and microtubules to the re-orientation response of the cell under cyclic uniaxial stretch. We treated fibroblasts seeded on ECM coated substrates with cytochalasin-D, to inhibit actin polymerization, and nocodazole, to disrupt microtubules, and uniaxially stretched the constructs for either 3 or 6 hours using 5% and 10% amplitude at a frequency of 1 Hz [23]. We also quantified and compared the SF and nuclear orientations between the different groups. These experiments clearly demonstrate that inhibition of microtubule polymerization did not abrogate the reorientation response of cells; in contrast, inhibition of actin led to complete loss of the reorientation response of the cells (Table 2; Supplementary Fig. S3). Further, actin and microtubule depolymerizations significantly abrogated the nuclear reorientation under cyclic stretch. Cytochalasin-D treated cells had ~7° change in nuclear orientation under high stretch amplitude and duration (A10T6) as compared to nocodazole treated cells which did not show any change in nuclear angular orientation under stretch (Fig. 2c). These results show conclusively the primary contribution of actin to the cellular reorientation responses as compared to that of the microtubules which play an important role in the nuclear reorientation.

**Table 2.**
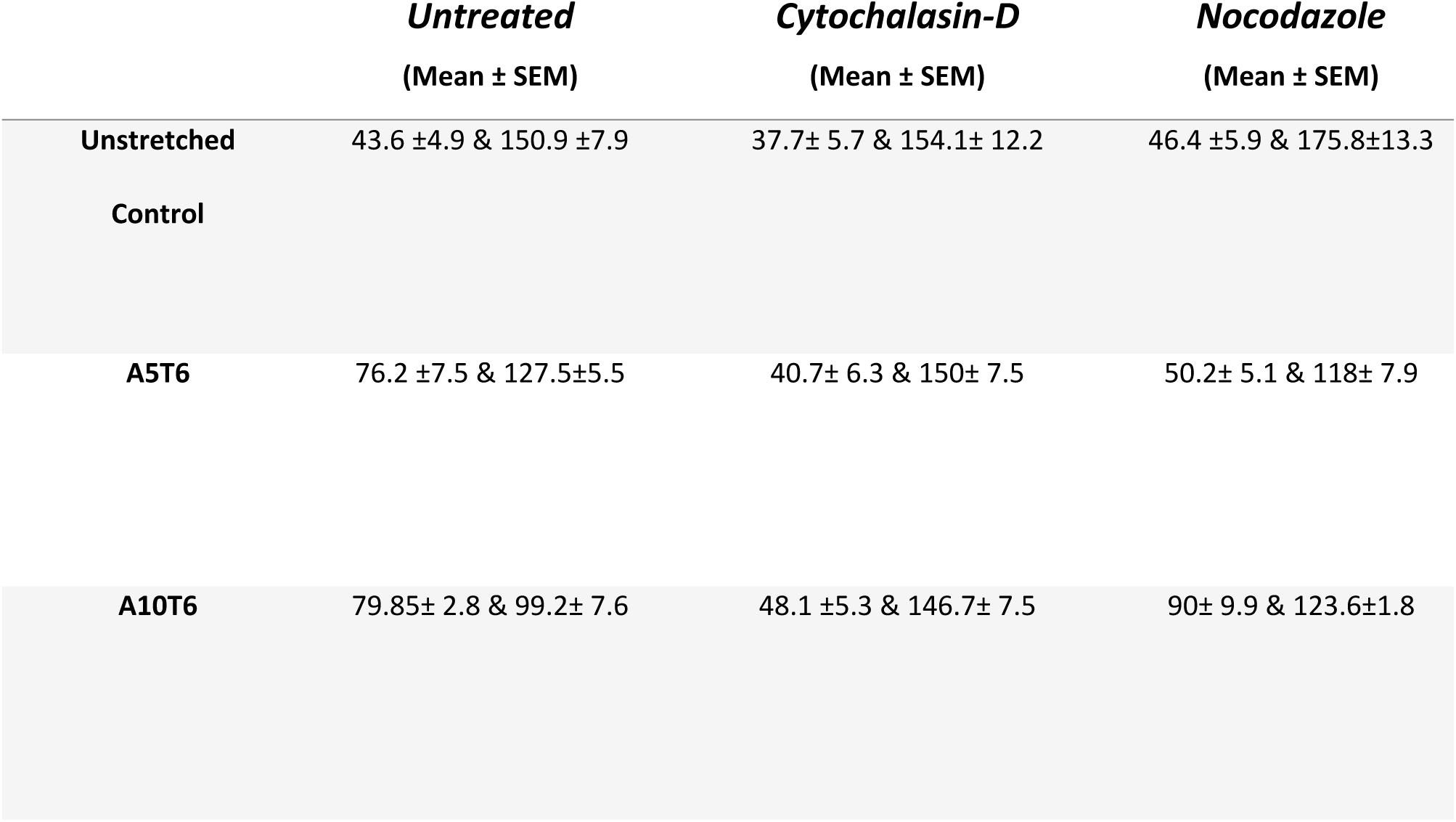
The effect of cytoskeletal inhibitors (cytochalasin-D and nocodazole) on the cells under uniaxial cyclic stretch were assessed using the SF angular orientation (Mean ± SEM) for fibroblasts cultured on PDMS membranes.

We quantified the differences in material properties of the cells before and after cyclic uniaxial stretch using an AFM (Atomic Force Microscope). Individual cells were indented with a spherical indenter with tip radius, *R_tip_*, and the modulus (E) was calculated using a thin layer, finite thickness, Hertz model (22,24) (**Supplementary information-1**). The force (F)-displacement (δ) relationship is given by

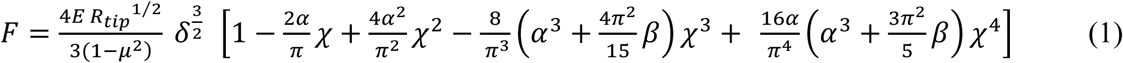

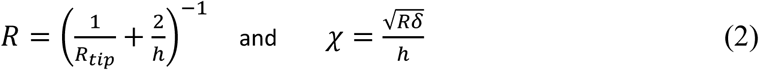

where *h* is the cell height and µ is the Poisson’s ratio of the cell (=0.5). α and β are constants that are functions of µ. All indentation experiments were performed for a constant indentation depth of 1 μm which corresponded to ~14% of the total cell height. Fibroblasts stretched at 10% for 6 hours (A10T6) were indented in the presence or absence of cytoskeletal inhibitors and the corresponding force-displacement curves were compared with unstretched controls. Our results show that cyclically stretched cells had significantly higher stiffness as compared to unstretched controls (p<.05; n~25 in each group; Fig. 2f). There was no change in the effective elastic modulus for cytochalasin-D treated cells under cyclic stretch. In contrast, nocodozole treated cells showed a significant increase in cell stiffness following cyclic stretch (p<.01). The cytoskeletal reorientation response in cells demonstrates the dominant contribution of SF in cell realignment under cyclic stretch. Depolymerization of microtubules, using nocodozole treatment, did not affect the stretch induced increased the cell stiffness and was linked to formation of SF which drives the cellular reorientation dynamics under cyclic stretch.

Results from cells stretched at 10% amplitude for 6 hours (A10T6) showed ~40% increase in the total actin fluorescence intensity as compared to unstretched controls (p<0.05) (Fig. 3f). These results on cyclically stretched cells were accompanied with increased SF lengths and higher cell stiffness measured using AFM. We measured intensity distributions corresponding to the F-actin fluorescence along the transverse sections of the cell at various points along the major axis of the cell to explore possible contributions of the cortical actin to the overall increase in cell stiffness due to cyclic stretching (Fig. 3a, b, c). The full width at half maximum (FWHM) values corresponding to the intensity distribution for each cell and the peak intensity values were used to characterize the thickness of the cortical actin for the cell (Fig. 3d, e). These results showed that the FWHM was 40% higher for cyclically stretched (A10T6) cells as compared to un-stretched controls (p<0.05); cyclic stretched cells also had significantly higher peak intensities as compared to control cells (p<0.01). Reinforcement of the cortical actin and an increase in the SF lengths in individual cells caused by cyclic uniaxial stretching hence resulted in the increased cell stiffness.

**Figure 3.**
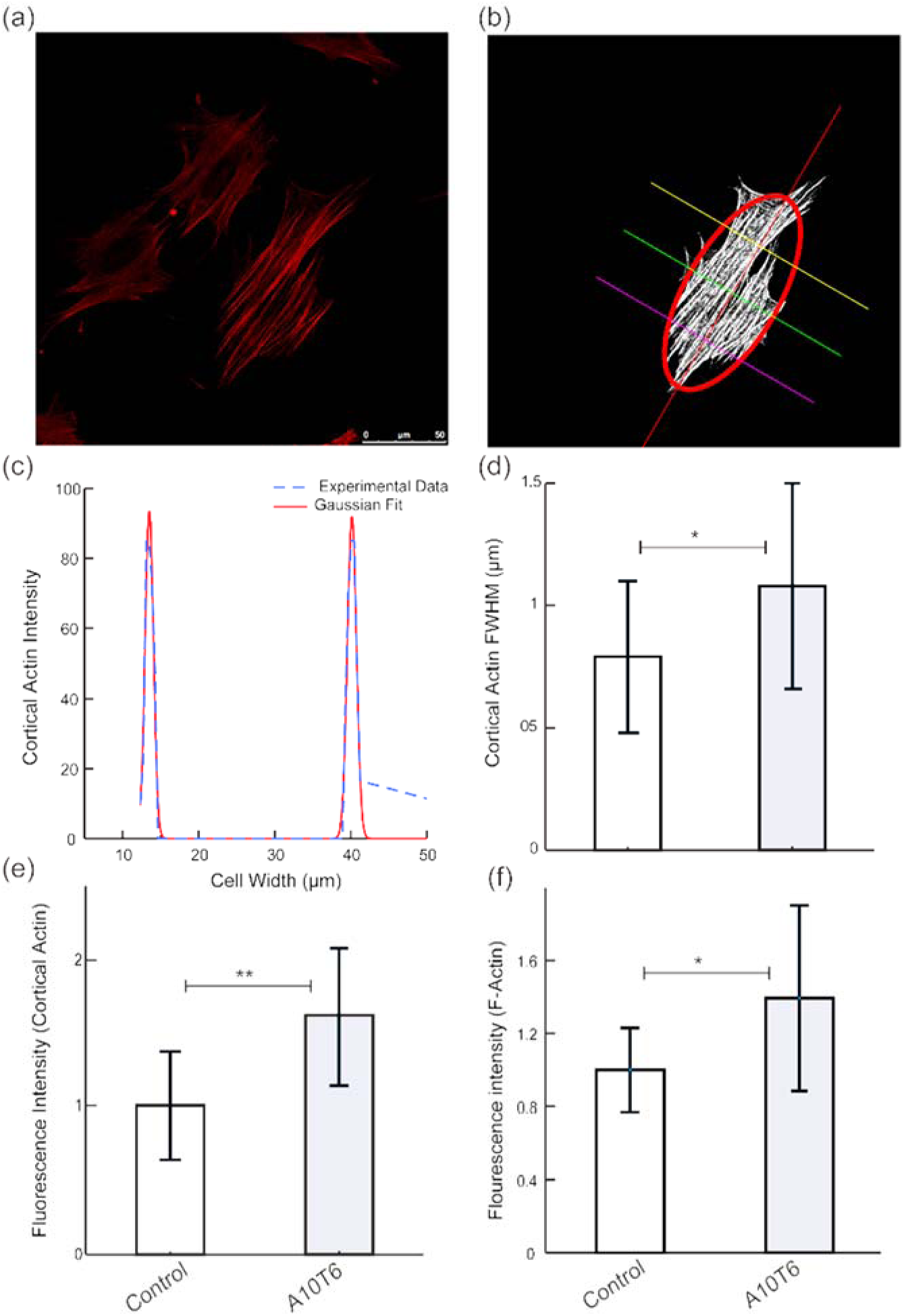
**a.** Confocal image of fibroblast stained with phalloidin to visualize actin. **b**. The corresponding binarized image of the cell was fit to an ellipse and the intensity distributions were calculated at three equidistant transverse sections along the major cell axis. **c.** A representative image of cortical actin intensity, calculated at one of the transverse sections along major axis, is shown **d**. The full width at half maxima (FWHM) was calculated to compute the actin cortex thickness **e.** The peak intensity of cortical actin in cells subjected to uniaxial cyclic stretch increased under stretch (A10T6) **f.** The total actin fluorescence intensity in cyclically stretched cells (A10T6) also increased as compared to unstretched controls. Significant differences in comparisons between the different groups are indicated for p<0.01 (**) and p<0.05 (*).

### Morphoelastic model of the cell under uniaxial cyclic stretch

Volumetric growth in the cell, associated with mass addition, is described within a finite strain elasticity framework based on deformations caused by non-uniform changes in the mass at the local level at each material point of the body [25]. The SF elongation in the fibroblast under cyclic stretch is described using the growth tensor, **G**, which produces a virtual stress-free and non-integrable configuration in the cell to generate residual stress. SF growth kinematics is accompanied with an elastic deformation, **A**, which is induced by growth and is essential in restoring compatibility of the overall deformation gradient, **F**, during growth. The elastic deformation tensor plays an important role in maintaining the cellular integrity and is related to **F** as:

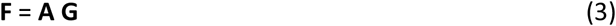

Coupling between the elastic responses and biological growth is integral in morphoelasticity and is consistent from kinematic and thermodynamic perspectives. Results from experiments to quantify the growth of SF in fibroblasts under uniaxial cyclic stretch were used to determine the growth kinematics based on the evolving configurations during cell reorientation under stretch (**Supplementary Information-2**).

We model the cell as a fiber-reinforced orthotropic hyperelastic material, comprised of two families of SF, based on the microscopy images (Fig. 1) [26,27]. A representative element of the cell is characterized through dependence on the elastic deformation, **A**, and is given by the strain energy function, *W(**A**)*. We assume that the contributions to *W* are given in terms of an isotropic component, ***W_iso_***, given by a neo-Hookean term related to the cytosol and an anisotropic component due to presence of two families of SF, ***W_aniso_*** which is given using a standard reinforcing solid model as

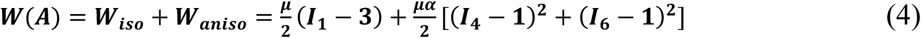

*μ* is the shear modulus (*μ* > 0) and *α* is a measurement of the additional fiber reinforcement (*α* > 0) [28]. The elastic counterpart of the right Cauchy-Green tensor ***C**^E^* = ***A^T^A***. *I_k_*(***A***); *k* = 1,4,6 are invariants of the elastic deformation given by [29]

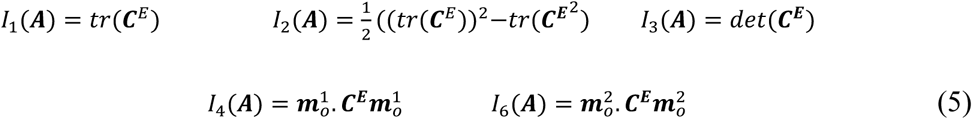

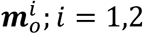 represents vectors corresponding to each of the two SF directions in the natural configuration in the absence of stretch.

We use experimental data on SF lengthening, measured at different amplitudes and for various amount of stretch durations, to formulate the evolution equation for increase in length (γ) of SF under cyclic stretch which is assumed to be a function of the applied stretch, *λ*, and time, t, and is given by

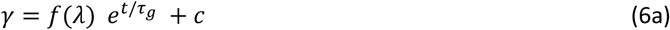

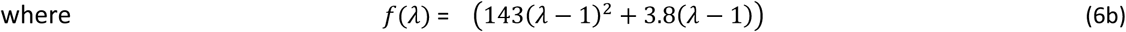

c is a constant with dimensions of length and is defined as the SF length in unstretched (control) cells; *τ_g_* is a growth time scale for the SF. The coefficients in *f*(*λ*) were obtained by fitting the experimental data to equation (6). We use these evolution equations and write the growth tensor for a single SF as

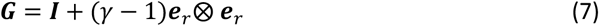

***e**_r_* represents the unit vector in the direction of SF growth. We next include thermodynamic restrictions for active growth based on the Claussius–Duhem inequality using the standard Coleman– Noll procedure (**Supplementary Information-3,4**).

We assume a form for the SF re-orientation equations along a uniform direction perpendicular to applied cyclic uniaxial stretch as

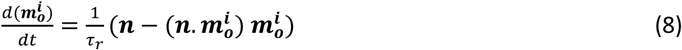

We have assumed the direction of applied uniaxial cyclic stretch is along the X-axis. The vector **n** represents the unit vector along the direction of preferred cytoskeletal orientation orthogonal to the applied stretch in the Y-axis (**e**_2_) [28,30]. Equation (8) implies that the two families of SF reorient perpendicular to the cyclic stretch direction over time and results in reinforcement. *τ_r_* is the remodeling time scale which depends on the stretch amplitude. The elongation and reorientation dynamics of SF in cells for 10% stretch amplitude over 6 hours (A10T6) is shown in **Supplementary Video 1**. The model provides insights into active SF growth that is accompanied with passive SF re-orientational dynamics.

### Model verification using uniaxial stretch data on cells

We tested the model predictions for changes in SF lengths and angles under uniaxial stretch and related their influence on the measured cell stiffness using experimental data. The evolution equation (6) for increase in length of SF under stretch is in agreement with the experimentally measured observations (r^2^ ~0.99). (Fig. 4a). The time scale for growth of SF in the model is described using *τ_g_* which is a function of the SF properties. We explored changes in SF growth by parametrically varying *τ_g_* in the model (Fig. 4c, d) for the 5% and 10% stretch amplitude and compared these results with experimental data; the model predictions match with experimental data for *τ_g_* =2 hours. The growth law also satisfies the thermodynamic incompatibility condition for active biological growth based on various possible fiber angles and experimental duration for different stretch amplitudes used in our study (**Supplementary information 4**). Fig. 4b shows the corresponding surface plot for the inequality term 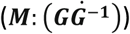 for 5% stretch amplitude as a function of time as well as fiber angles. ***M*** is the Mandel stress tensor which is a thermodynamic conjugate to the velocity gradient, 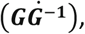 satisfies frame indifference, and is used as a natural descriptor for growth processes [31,32]. From Fig. 3b, we note that 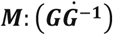 is positive for all possible values of time and fiber angle in our model which satisfies the thermodynamic constraints.

**Figure 4.**
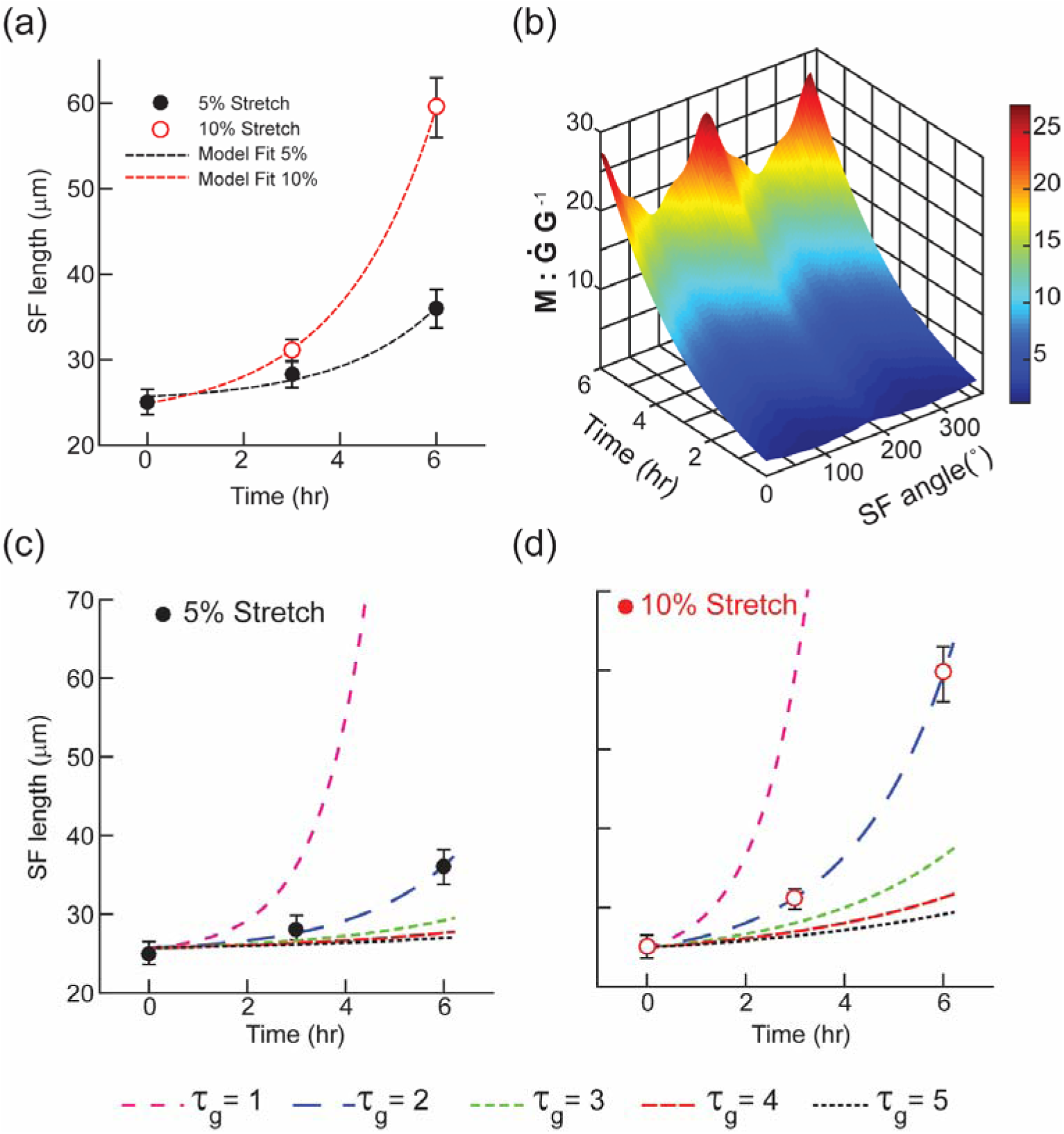
**a**. Changes in SF lengths under cyclic stretch (5% amplitude and 10% amplitude) were calculated from experimental values (mean ± SEM) and were used to obtain the growth evolution law for SF (equation (6)). The unknown constants in the equation were obtained by fitting the experimental data to equation (6) (r^2^ = 0.99). **b.** Thermodynamic inequality for active growth process was assessed as a function of experiment duration and SF orientation. **c.** & **d**. The growth time scale (*τ_g_*) was parametrically varied in the SF growth model for 5% stretch (black) and 10% stretch (red). Experimental results are also plotted on the figures using black and solid circles for 5% and 10% amplitude.

The reorientation law predicts SF orientation under various magnitudes of stretch and time duration using inputs corresponding to the mean SF angular distributions. We compared model predictions from fiber reorientation law with the experimentally obtained SF orientation distributions that were quantified using confocal images for both 5% and 10% cyclic stretch data. The model shows a good match with experimental data for increased SF orientational dynamics with higher stretch (Fig. 5a, b). Remodeling time scale for SF orientation, *τ_r_*, increased significantly with the magnitude of applied stretch. Experimental results are superposed on the model predictions; these demonstrate a good match for *τ_r_* =3 hours for 10% stretch and *τ_r_* =9 hours for 5% stretch (r^2^>0.9 for both). That is, cells reorient to the orthogonal direction to direction of stretch in a shorter time when subjected to higher stretches. Supplementary Fig. S4 shows parametric variations corresponding to various initial conditions and times for the SF reorientation law.

**Figure 5.**
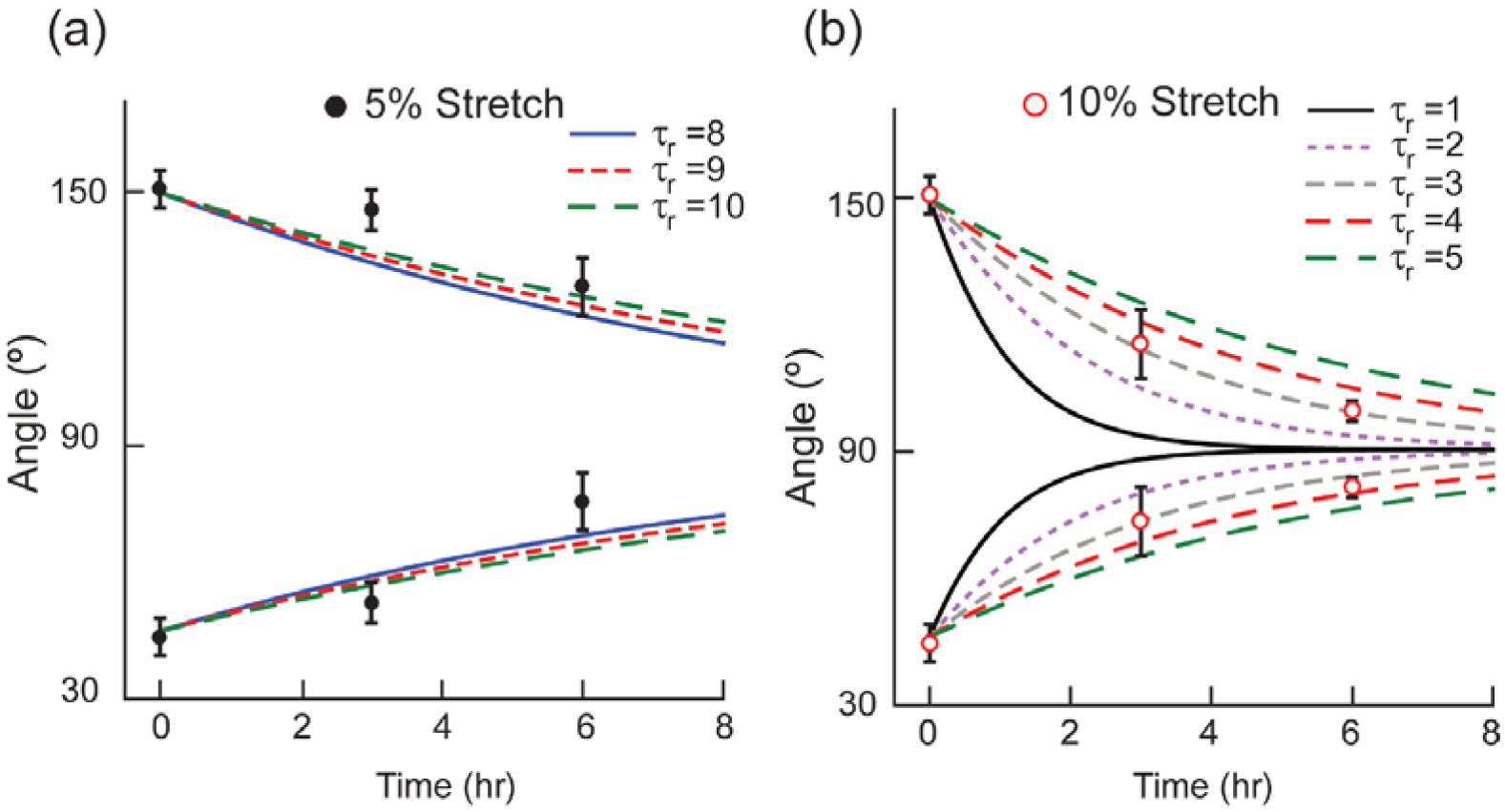
**a&b**. The computed SF reorientation under cyclic uniaxial stretch agreed with experimental observations which demonstrates the importance of SF remodeling time scale (*τ_r_*) as a function of the stretch amplitude. *τ_r_* was 9 hours for 5% and 3 hrs for 10% stretch respectively (r^2^ > 0.9 for each)

We calculated changes in the effective elastic modulus, E_eff_, of cells as function of fiber reorientation angle and time (equations (8). E_eff_ in a given direction is defined as the gradient of uniaxial tension, **N**, with respect to the stretch in that direction, is given in terms of the stress-free configuration as 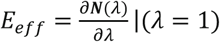 (31) (**Supplementary Information-6**). Variations in E_eff_ with fiber angles (*θ*_1_ ∈ (0°, 90°) *and θ*_2_ ∈ (90°, 180°)) from the two SF families are shown in Fig. 6a, b. These plots show an increase in *E_eff_* when both families of fibers align perpendicular to the applied stretch direction. Variations in *E_eff_* with time and the stress distributions with variations in fiber angles of two SF family (*θ*_1_ *and θ*_2_) for 5% and 10% stretch amplitude were also calculated (Fig. 6c, 6d). Stresses obtained from experimentally obtained fiber distributions at various stretch amplitude and time points using structure tensors based on fiber orientation distribution functions (**Supplementary Information-7**) show a similar trend as that predicted by the model for reorientation under cyclic stretch along the minimum stress direction (Fig. 6d). These observations are in good agreement with results from by De *et al* (33) who suggested that cellular reorientation under uniaxial cyclic stretch occured along the minimal matrix stress direction.

**Figure 6.**
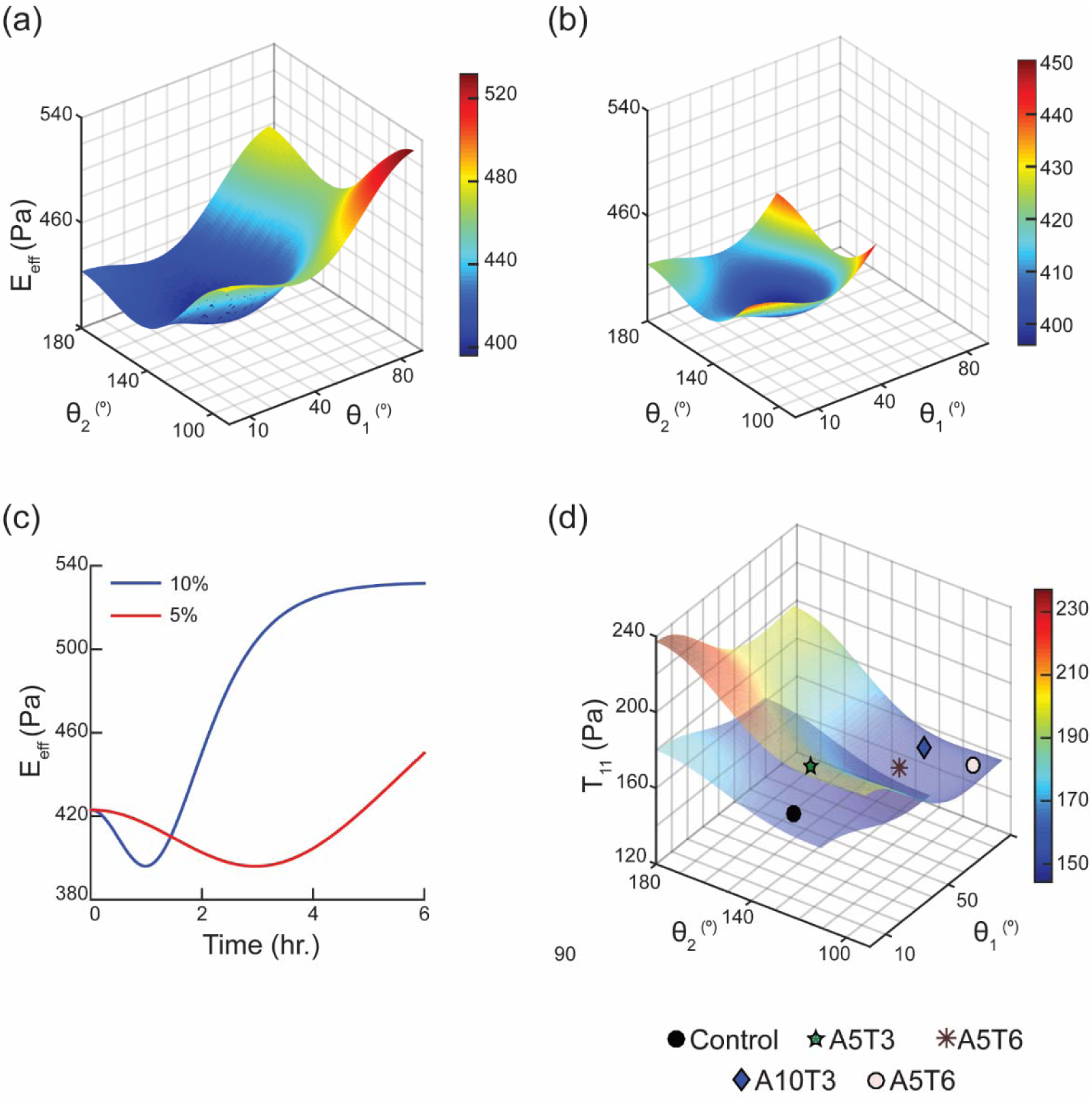
**a&b**. Changes to the effective elastic modulus, E_eff_, of the cell (in Pa), subjected to 5% and 10% amplitude uniaxial cyclic stretch, were measured and are plotted as a function SF alignment angle. **c**. Changes in E_eff_ based on the model are shown as a function of SF reorientation under cyclic stretch (5% stretch in red and 10% stretch shown in blue). **d**. Changes in cell tractions (T_11_) under uniaxial stretch from the model calculations are compared with the experimental results.

## Discussion

Growth and dynamics of SF networks are important in maintaining cell shape, control of cellular mechanical properties and mechanotransduction processes underlying the adhesion and migration of cells. The primary aim of this study was to quantify the morphomechanical response of fibroblasts under uniaxial cyclic stretch. There are three main implications of this work. First, we show that cyclic stretch induced SF growth and reorientation alters the cellular morphometrics. SF elongate based on the amplitude of cyclic stretch; their orientations changed from an initial random configuration in unstretched cells to a uniform direction perpendicular to the applied stretch. Cortical actin thickness increased with applied stretch which resulted in an increase in the effective elastic modulus of the cell measured using AFM. Second, we quantified the differential roles of actin and microtubules in cellular and nuclear reorientations under stretch. We show that actin is essential to the cellular orientation changes under cyclic stretch and contributes to the measured cell stiffness; in contrast, microtubules influence nuclear reorientations. Third, we use a nonlinear, orthotropic, hyperelastic, and fiber-reinforced constitutive model for the cell to elucidate the combined effects of amplitude and duration of uniaxial cyclic stretch using a growth and remodeling framework. The model uses SF evolution equations that depend on the stretch amplitude and time and were determined experimentally to predict the effects of uniaxial cyclic stretch on SF dynamics.

Actin growth dynamics is exquisitely sensitive to mechanical stimuli [34,35]. Dense branched actin networks grown on functionalized surfaces demonstrate nonlinear mechanical responses [34]. Parekh *et al* [36] measured the force-velocity relationships of growing actin networks *in-vitro* using a differential AFM and showed the dependence of actin network growth velocity on the history of applied loading. Mechanical stimuli like torsion and fluctuations in applied loading also influence the growth of actin networks [37]. These studies were however reported on *in vitro* preparations of isolated and purified actin networks; our results show similar responses for actin networks in fibroblast cells under stretch. The resulting changes to the material properties of the cells, through a process of *morphoelasticity*, has not been demonstrated earlier at the cellular level. The actin cortex also plays an important role in maintaining cellular shape [38]. We show that the application of uniaxial cyclic stretch increases cortical actin thickness which reinforces the cytoskeleton and also results in an increase in the cell stiffness (Fig. 2f; Fig 3d). Hasse and Pelling showed the importance of actin cortex in controlling cellular deformation by applying precise nano Newton forces [39]. Comparison of the cell modulus to the actin network orientations based on experimental data and model predictions show that cells with higher SF dispersions have a more compliant response as compared cells with highly aligned SFs (Fig. 2f; Fig 3d) [22,40].

Uniaxial cyclic stretch experiments in our study show that the growth realignment of SF is dependent on the amplitude of applied external stretch; SF aligned from a random configuration to a uniform direction perpendicular to the direction of cyclic stretch (Fig. 1; Table 1). The hyperelastic model for the cell under external cyclic stretch is based on the assumption of the SF as an elastic structure that undergo elongation and realignment. The persistence length for a SF, *ξ*_p_, given as a ratio of the bending stiffness to statistical fluctuations, is typically greater than 50 µm [41,42]. Filaments with lengths smaller than *ξ*_p_ are flexible and those with greater dimensions are generally described using statistical methods to include the effects of thermal fluctuations. The longest SF length in the fibroblasts in our study was less than *ξ*_p_ which justified the use of elastic rod approaches to model SF growth dynamics.

We used a phenomenological nonlinear, stretch-dependent evolution equation for SF elongation that was dependent on time and cyclic stretch amplitude from the experimental results (Fig. 3). We used two different time scales in our model to represent changes in the SF length and orientation under cyclic uniaxial stretch. The time scale for fiber growth, *τ_g_*, is constant and depends on the intrinsic properties of the SF. The second time scale, *τ_r_*, represents SF remodeling time scale which is sensitive to applied stretch and decreases with an increase in stretching amplitude (Fig. 4). We quantified the individual contributions of SF and microtubules in cellular and nuclear reorientation under stretch and show that microtubule depolymerization did not significantly affect cellular reorientation response. However, actin depolymerisation resulted in almost complete loss of the cellular reorientation response (Table 2; Supplementary Fig. S3). These results are in close agreement with a previous study by Goldyn *et al* (17) who showed dependence of force-induced cell reorientation on FA sliding alone and was largely independent of the contributions from microtubule networks. AFM indentation experiments in our study also show that the depolymerization of microtubules did not change the cellular stiffness under stretch; formation and re-organization of SFs in response to stretch induced cytoskeletal stiffening.

Stretch induced cytoskeletal reorganization has previously been linked to stress or strain homeostasis in the cells [33,43]. Recent studies show that the direction of cell realignment under the application of stretch depended on the traction boundary conditions [20]. The reorientation dynamics of individual cells under uniaxial cyclic stretch resulted from a minimization of the elastic strain energy in the cell [4]. We used a morphoelasticity framework to show that the cyclic stretch dependent cellular morpho-mechanical properties are a result of SF growth and reorientation in a direction perpendicular to the cyclic stretch direction. These observations are in agreement with previously published works by De *et al* who predicted the direction of cellular reorientation along the minimal matrix stress direction [33]. One limitation in our model is we have not considered the effects of branched networks and have only included straight filaments. SF are thick and stable and generally distinguished into four categories as ventral and dorsal SF, transverse arcs, and perinuclear actin cap [44]. We have modelled the growth and reorientation dynamics of ventral SF alone and show the importance of Sf reinforcement in the measured changes to the mechanical properties of the cell.

To conclude, we used a novel growth and remodeling framework to study the effects of uniaxial cyclic stretch in fibroblasts using experimental stretching methods in combination with a continuum mechanics framework. The model uses experimentally motivated evolution equations for SF growth and remodeling that have not been reported earlier. We show an increase in the cellular stiffness under cyclic stretch which is linked to SF growth and reorientation in a direction orthogonal to the direction of stretch. These studies show the importance of uniaxial stretching in mechanotransduction processes related to changes in the cytoskeleton which are essential to model diseases like aneurysm growth and fibrosis that currently use qualitative representation of the cell under stretch.

## MATERIALS AND METHODS

### 1. Design and assembly of the bioreactor for cell stretching

The custom designed cell stretcher allows for both uniaxial and biaxial stretching with equal and different strain rates on each actuator such that the central point for viewing is fixed. We designed a novel clamping arrangement to clamp a thin stretchable membrane (width 10mm x length 20mm x thickness 150 μm) between two opposite translating actuator arms for the dynamic stretch experiments. The clamps were 3D printed using a biocompatible acrylic compound (Verro White; Stratasys) (Supplementary Fig. 1a, b). The cell stretcher was operated using user-controlled stretch amplitude and frequency inputs over extended time periods and placed within a humidified temperature-controlled incubator to maintain sterility.

### 2. Strain quantification

Strain quantification experiments were performed to calibrate the novel clamping design for the specified stretch amplitude during cyclic stretch experiments. In-plane strains were quantified by tracking fiduciary markers in the PDMS (poly di-methyl siloxane; Sylgard ®184, Dow Corning) sheet, mixed in the ratio of 10:1 elastomer to curing agent, using an overhead video camera (Sony HD HDR-XR100E). Marker centroids were obtained using segmentation and the corresponding displacements were quantified. Marker displacements from the referential to the deformed configuration were measured and the deformation gradient, **F**, was computed as ***F = I + H***, where ***H**=∂**u**/∂**X*** is the displacement gradient in the referential configuration. A strain interpolation algorithm, implemented using MATLAB (v R2016a), was used to quantify in-plane Green-Lagrange strains in the specimen using measured values of the deformation gradients [27, 48]. These data were used to define a central homogeneous region, where strains were constant in the sample, and used to quantify cell behaviours.

### 3. Cell culture and immunofluorescence staining

NIH 3T3 fibroblast cells were cultured in Dulbecco’s minimum essential medium (DMEM; Sigma Aldrich) supplemented with 10% fetal bovine serum (GIBCO BRL) and 1 vol% of penicillin-streptomycin (Sigma Aldrich) in a humidified temperature-controlled incubator (Panasonic Inc.) at 37°C and 5% CO_2_. The same medium was also used during cyclic stretching experiments performed using the custom bioreactor within the incubator. Following stretching experiments, cells were fixed in 4% paraformaldehyde, permeabilized in 0.5% TritonX-100 (Sigma) solution for 10 minutes and stained either for actin using rhodamine phalloidin (Thermofisher 1:200) or microtubules using α-tubulin FITC (Invitrogen 1:200), and a nuclear DAPI counterstain (Thermofisher 1:400). Samples were imaged using a confocal microscope (Leica Microsystems, TCS SP5 II) using 10X, and 63X oil immersion objectives to visualize the cytoskeletal organization.

### 4. Uniaxial cyclic stretching of fibroblasts in the bioreactor and cytoskeletal inhibitor treatments

Thin flexible PDMS membranes were used as substrates to culture cells. PDMS was mixed in the ratio of 10:1 elastomer to curing agent, degassed in a vacuum pump, and spin coated on triacetylene treated glass slides at 300 rpm and 60 seconds to yield 150 μm thick sheets. The substrates were plasma treated for 2 minutes, incubated with 30μg/ml of fibronectin solution at 37°C for 45 minutes, and washed twice with PBS. Prepared substrates were incubated with DMEM containing 10% FBS for 30 minutes followed by seeding of cells. Cell seeded constructs were incubated overnight at 37°C and 5% CO_2_ to allow cell attachment and were stretched cyclically (5% and 10% amplitude) at 1 Hz for varied durations to explore the temporal dependence of stretching on cell alignment (3 hours and 6 hours).

To delineate the individual contributions of actin and microtubule in cellular response under cyclic stretch, we treated cells with either 0.5 µM concentration of cytochalasin-D (Sigma-Aldrich; USA) to depolymerize actin or 0.5 µM concentration nocodazole (Sigma-Aldrich) to disrupt microtubules [23]. We ensured that the same concentration of the drug was present during the stretching experiment to prevent possible repolymerizations of actin and microtubules that may contribute to possible artefacts in the experiments.

### 5. Quantification of SF orientation under cyclic stretch

Confocal images of cells stained for actin were analysed using a custom MATLAB program (Mathworks, 2016b) [21,22]. In this procedure, a binarized image of the actin intensity in the cell was obtained and represented using a distribution function f(x,y); (x,y) defines a point in the 2-D image. We calculated the Fast Fourier transformation (FFT) of this image as 𝓕 *(f (x, y)) =(u,v)* where (u,v) denotes a point in the Fourier space. The power spectrum of this matrix was obtained by *P (u, v) =F (u, v). F* (u, v)*, F* is the complex conjugate of the function, F in this expression. The SF were distinguished based on the spatial frequencies and orientations in Fourier space. A wedge-shaped orientation filter was used to quantify the SF orientation distributions (Supplementary Fig. S2) [21].

### 6. Measurement of effective modulus of the cell using Atomic Force Microscope (AFM)

We used a Park Systems XE-Bio AFM to quantify the effective modulus of the cell using an App Nano Hydra 6V-200NG-TL silicon dioxide cantilever with an attached spherical bead of diameter 5.2 μm. The stiffness of the cantilever based on manufacturer sheet was 0.045 N/m and was determined to be 0.041 N/m using thermal tuning method. Specific locations for indentations were selected using the cell topography to be approximately equidistant from the nucleus and the cell edge to minimize the influence of the nucleus. Indentation experiments were performed for each cell using cells in Hank’s balanced salt solution and the corresponding force-displacement curves for each location were used to quantify the effective Young’s modulus of the cell using the material properties of the cantilever and a modified Hertzian contact mechanics model [24]. The force curve corresponding to the approach of the tip towards the substrate was measured and a constant indentation depth of 1 μm was maintained. For indentation experiments on cells treated with cytoskeletal depolymerizers, we ensured that the optimum concentration of the specific drug was present throughout the experiment.

## Supporting information

Temporal changes in stress fiber lengths and orientations for a fibroblast under 10% cyclic uniaxial stretch

## List of Supplementary Figures

**Figure S1.**
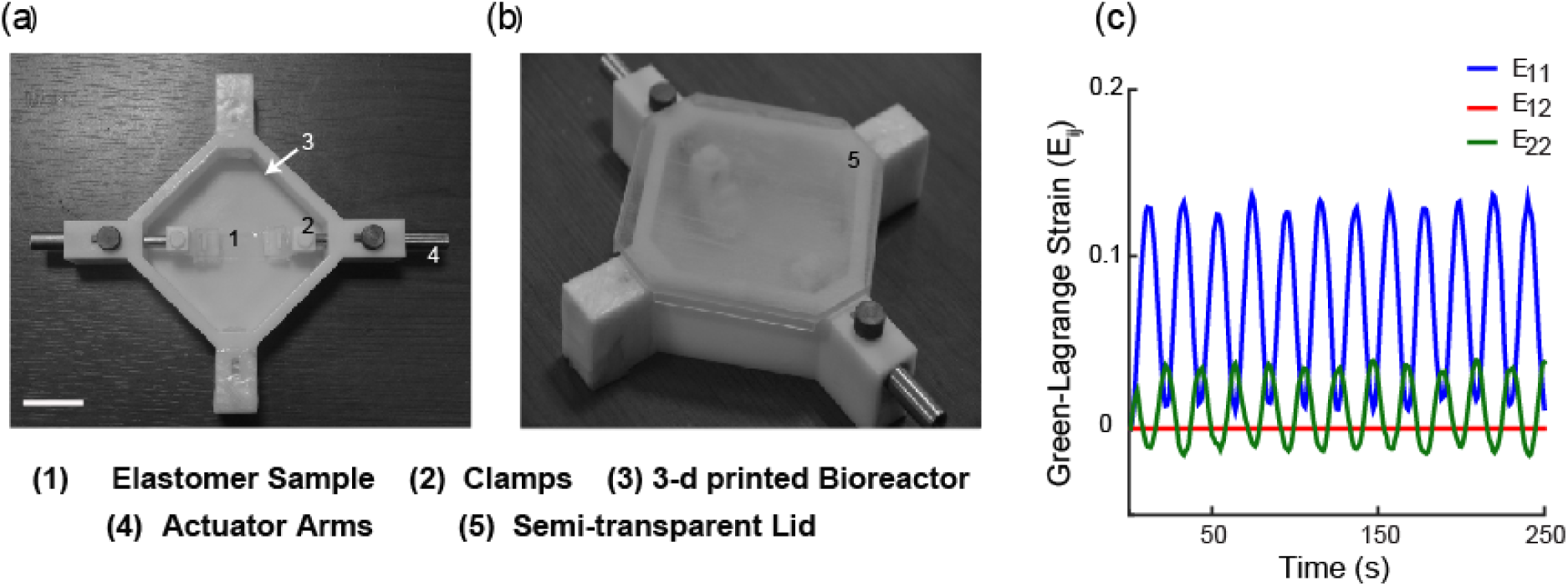
**a & b.** Custom designed cell stretcher and bioreactor. A thin PDMS membrane is held using custom designed clamps in the 3D printed bioreactor that may be subjected to uniaxial or biaxial stretching. **c** Quantification of Green-Lagrange strains under uniaxial cyclic strain (15% amplitude) are shown for the custom designed clamps using particle tracking algorithm using computed strain displacements based on the underlying markers on the substrate.

**Figure S2.**
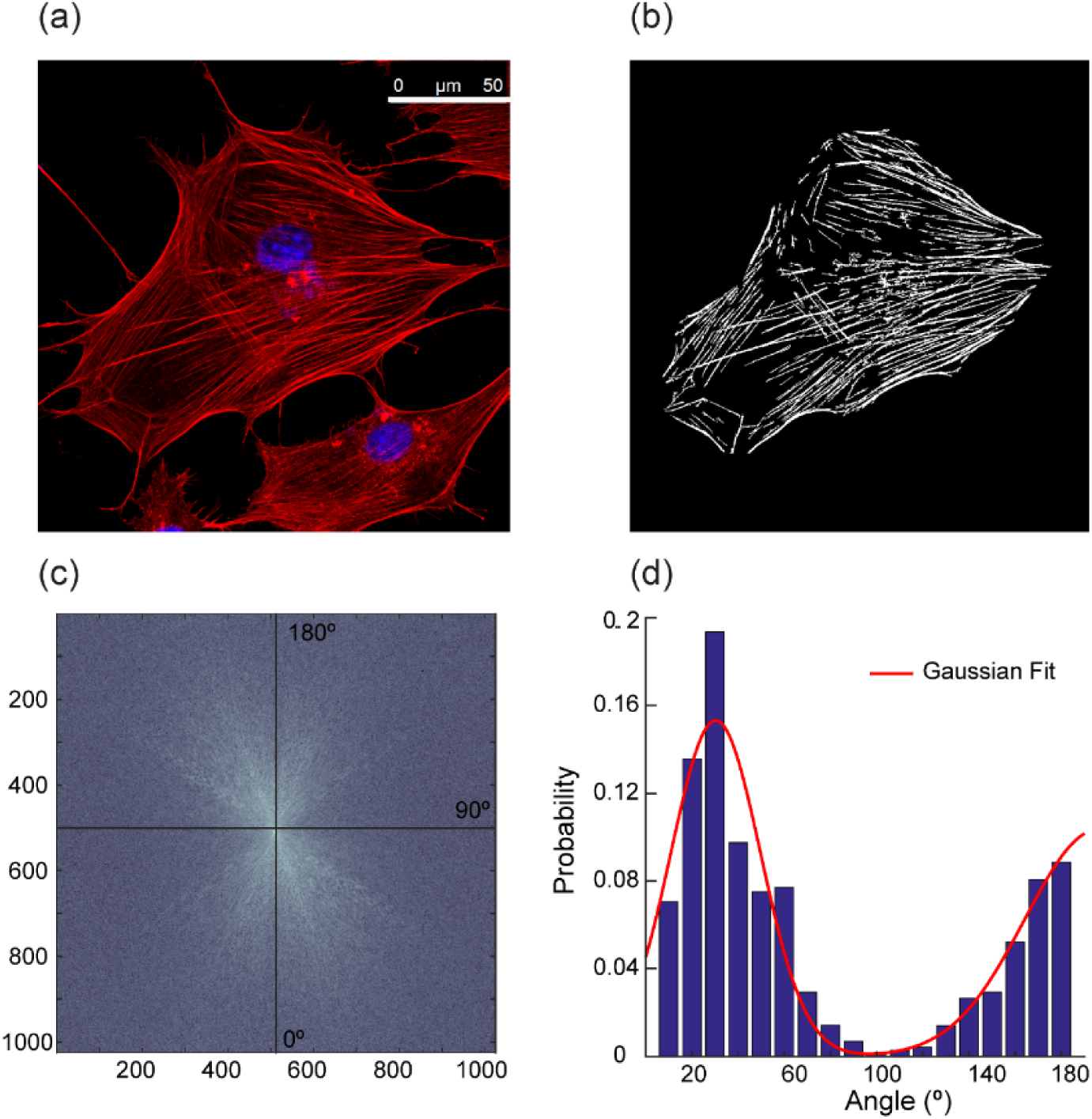
**a.** Confocal images show an individual 3T3 fibroblast cell, stained with phalloidin to visualize SF and DAPI to identify the nucleus. **b.** Corresponding binarized image of the confocal image is shown. **c.** The computed Fourier power spectrum was obtained using the binary image **d.** Distribution of SF in the cell was calculated using their spatial frequency distribution.

**Figure S3.**
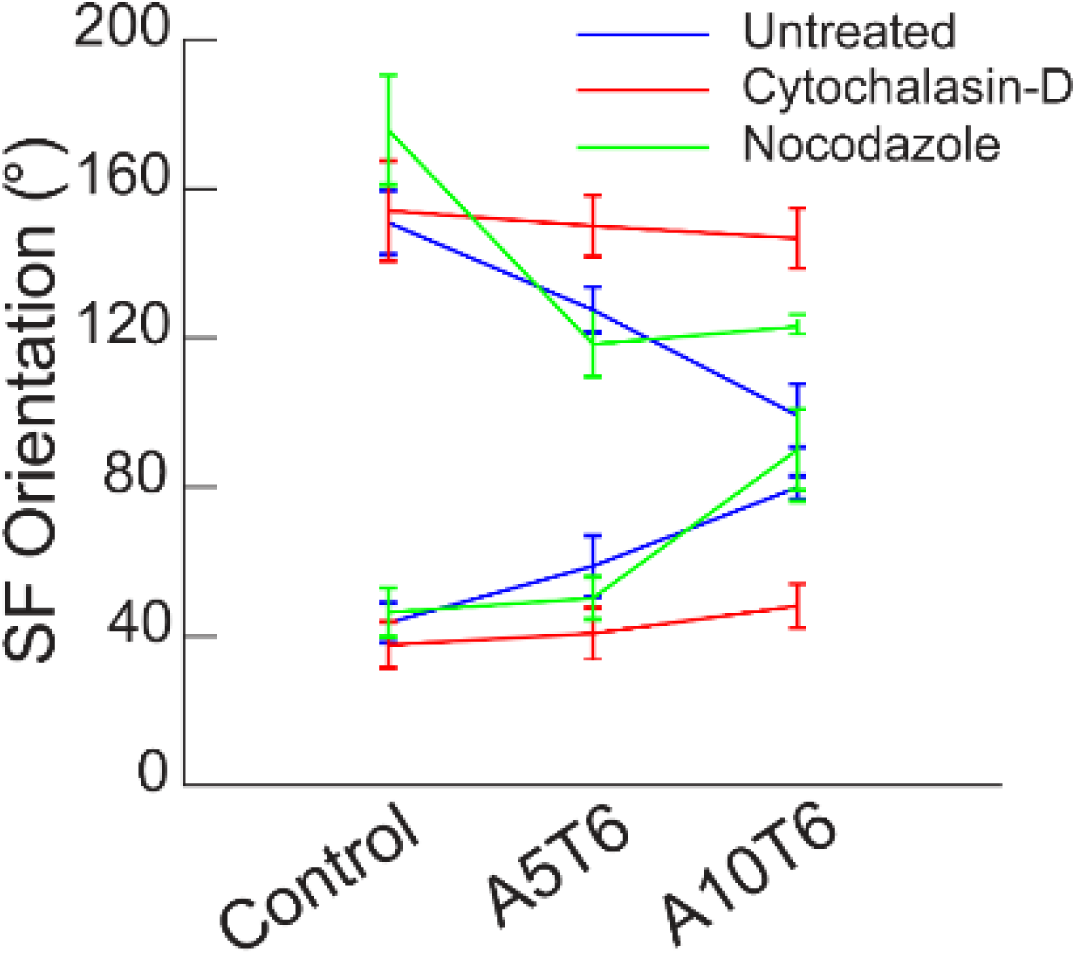
Angular orientation of the SF (Mean ± St. Dev) under unidirectional cyclic stretch (1 Hz) with and without cytoskeletal disruptors (cytochalasin-D and nocodazole)

**Figure S4.**
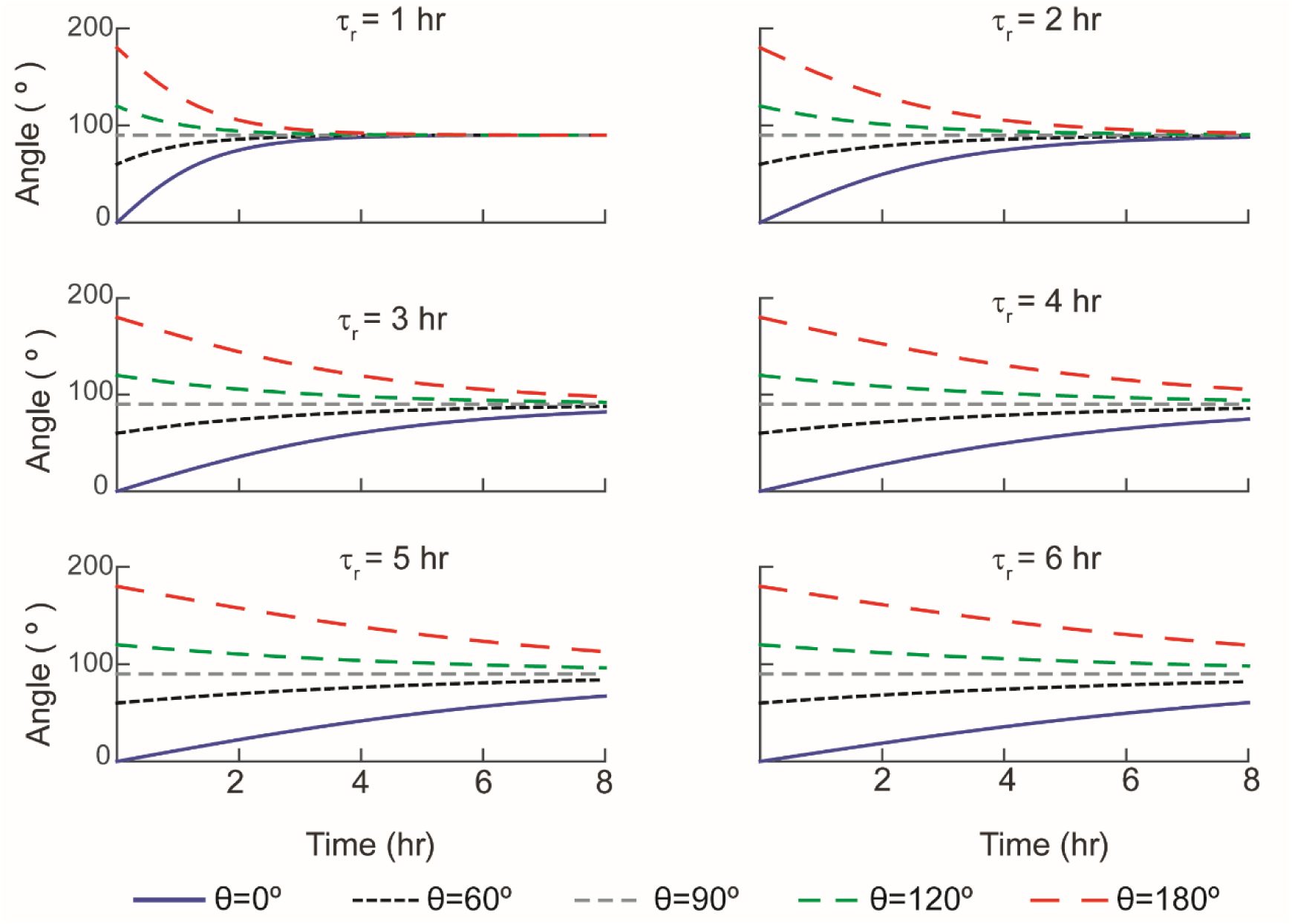
Parametric study of the SF reorientation model for various remodeling time scale (*τ_r_*) in hours.

## SUPPLEMENTARY INFORMATION

### 1. Thin layer, finite thickness Hertzian model to compute the effective elastic modulus

The force measured during indentation on the cell (F) is related to the indentation depth of the cantilever tip, δ, as [1]

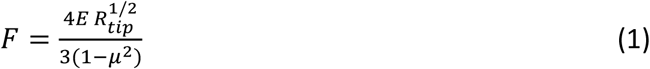

*µ* is the Poisson’s ratio, E is the Young’s modulus, and R_tip_ is the radius of the spherical cantilever tip. The cell was assumed to be incompressible with µ = 0.5. The tip-sample separation, Δ, is related to indentation depth, δ, by a constant, C, and is given by δ =C – Δ [2].

Using this expression, we modify the Hertzian relation as

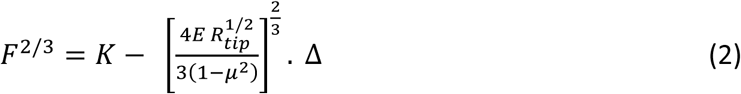

K is another constant and depends on both the coordinates of the substrate contact point with the cantilever tip and the geometry of the tip. *E* can be directly calculated as a scaled slope of the linear relation of the F^2/3^-Δ curve. The point of maximum slope change accompanied with change in sign of the force is considered to be the point of contact of the cantilever tip with the cell. The force data is divided into a region prior to the contact point of the cantilever tip with the cell and a region after contact. r^2^ values were calculated for each of the raw data curves to estimate the goodness of the linear fit; values with the mean r^2^ value higher than 0.9 are reported in this study.

The calculated values of E were normalized using a thin layer, finite thickness Hertz contact model to determine the elastic modulus of soft materials [3]. The force (F)-displacement (δ) relationship is given as

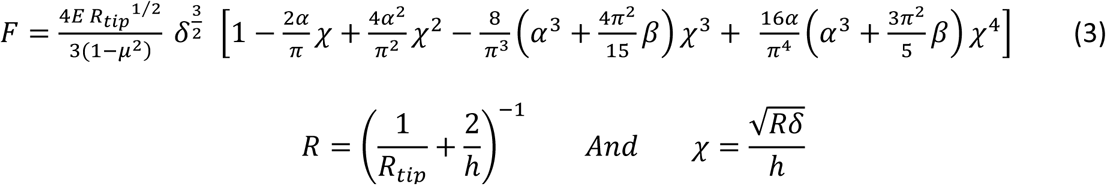

In this expression, *h* is cell height, µ is the Poisson’s ration and the constants α and β are functions of µ.

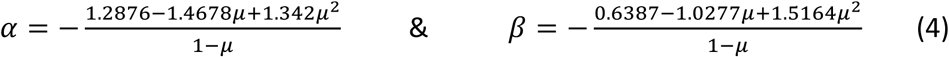

The heights of the cells were taken to be 7 μm based on Z-stacks of confocal images. For this analysis, δ = 1 μm (~14% of the total cell height) for all data which lies within the 4-30% range of the cell height reported in literature that are required for the equations to be used for such studies.

### 2. Constitutive modelling of fibroblast in a morphoelastic framework

We use a fiber-reinforced orthotropic hyperelastic material model to quantify the SF reorientation dynamics of fibroblasts under cyclic stretch. Our results show distinct growth and reorientation of the SF along a near perpendicular direction perpendicular to stretch. We also observe remodeling, in terms of increased cell stiffness and change in effective elastic modulus, in the fibroblasts following cyclic stretch. The strain energy density function of the cell combines a neo-Hookean response for the cytoplasm and a ‘*standard reinforcing model’* [4,5] for the two families of SF in the cell, as obtained from our experimental observations. The direction of fiber orientation is obtained from the angular distributions of SF from two term Gaussian distribution functions.

The deformation gradient, ***F***, is written in terms of a multiplicative decomposition of the growth tensor, ***G*** and the elastic deformation tensor, ***A***, in the framework of morphoelasticity [6].

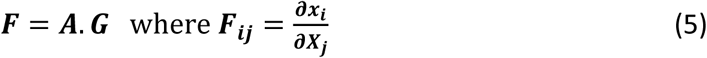

The strain energy density function of the cell is expressed as a function of the elastic deformation ***A,*** such that,

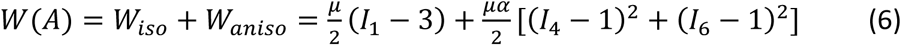

Where *μ* refers to the shear modulus (*μ* > 0) and *α* is a measurement of the additional fiber reinforcement (*α* > 0). The strain energy function also satisfies the condition [7]

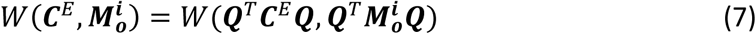

The structure tensor is given as 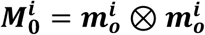, where 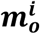 denotes the stress fiber distributions. ***C^E^ = A^T^A*** [8] and ***Q*** is an arbitrary orthogonal second-order tensor. The invariants used in the strain energy function are given as: -

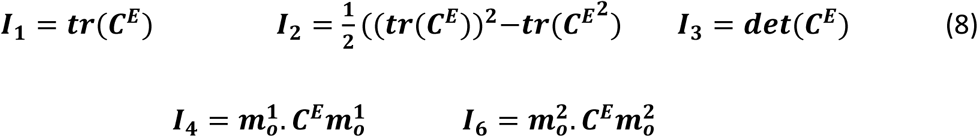

We consider the biological cell as an incompressible material, such that *det(**A**)* = 1. For uniaxial stretch, the elastic deformation tensor, ***A*** is given by

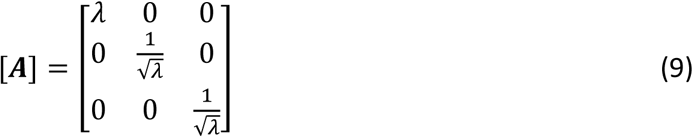

where *λ* is the magnitude of stretch applied along X-axis. The Cauchy stress can be computed as a function of the elastic deformation ***A***, using the relation [9]: -

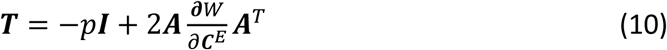

Where p refers to the Lagrange multiplier due to incompressibility assumption.

The condition for mechanical equilibrium is given by ***∇. T*** = 0. In our model, the uniaxial cyclic stretch is applied in the X-direction and the cell is adhered on the substrate in the X-Y plane.

Therefore, the unit vector along the direction SF is given as

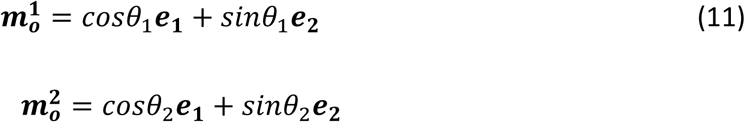

Using the above relations, we can compute the following identities

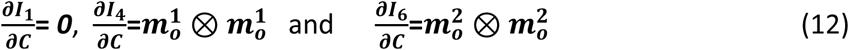

Substituting these expressions, the constitutive equation for the cell can be written as

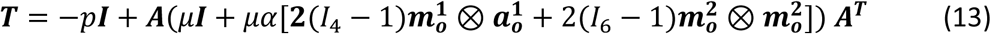

The corresponding tractions for the cell in the directions of ***e*_1_, *e*_2_** and ***e*_3_** are computed as

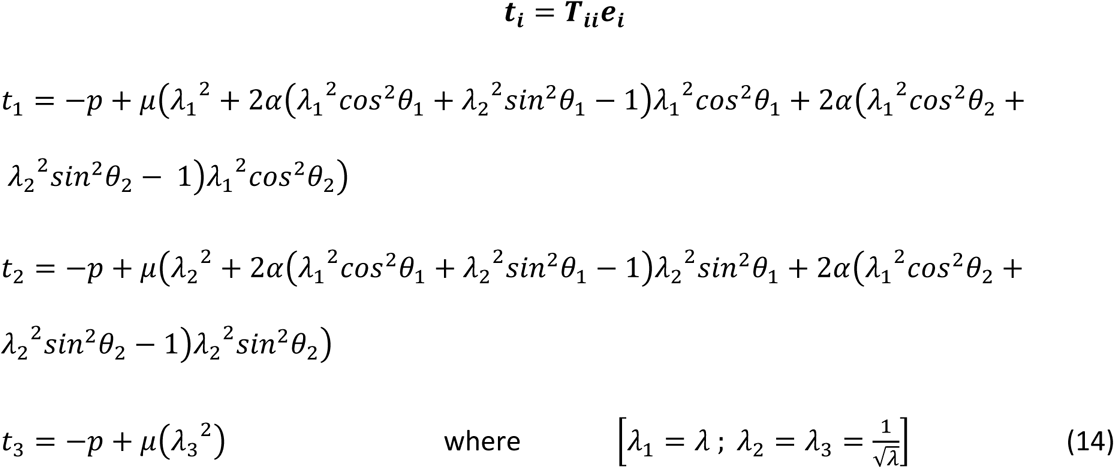

### 3. Growth Law for evolution in the SF length under cyclic uniaxial stretch

We propose a growth law for the SF under uniaxial cyclic stretch as a combined function of both the stretch amplitude and experimental duration.

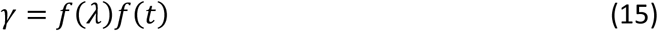

Based on our experimental observation, we formulate an evolution law for the increase in length of SF under cyclic stretch as

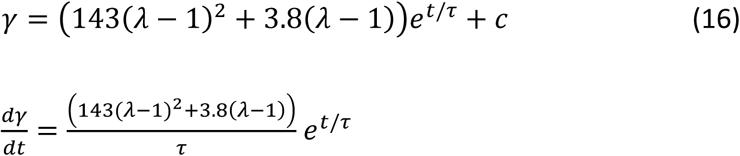

The parameters are obtained by curve-fitting to the experimental data, where c= is a constant indicative of the length of SF in unstretched cells, *τ* is equivalent to a growth time scale of the cell; *λ* is the corresponding stretch and ***t*** is time.

Based on the growth law of the SF under stretch, we now construct the growth tensor for the growth of a single actin stress fiber in the cylindrical co-ordinate system:

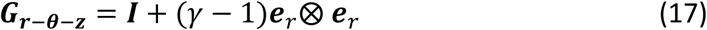

*γ* corresponds to the growth of actin SF in the radial direction. We can then transform *G_r-0-z_* from the cylindrical co-ordinate system to Cartesian using the transformation: -

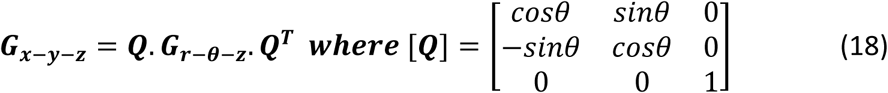

### 4. Thermodynamic Constraints on the Growth Law

Thermodynamic restrictions on the growth law were computed from the Claussius– Duhem inequality using the standard the *Coleman–Noll* procedure. Neglecting temperature gradients, the thermodynamic inequality for active growth processes is written as [10]: -

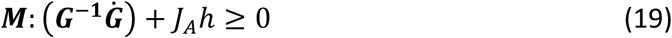

***M*** is the Mandel stress tensor; 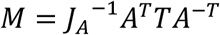 and the entropy contribution in the process is given by *h*. In our case *J*_A_=1 due to incompressibility. We consider the SF as a network non-Gaussian chain and use a statistical treatment of randomly jointed chains to quantify the change in entropy [11] under uniaxial tension 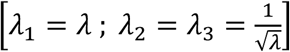

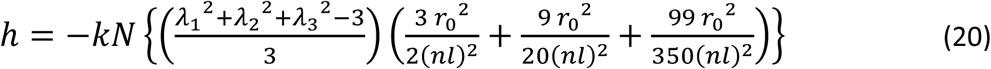

*n* is the number of links in one chain and *l* corresponds to the length of each link. The ratio 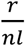 is a measure of the fractional extension of the chain; *r*_0_^2^ is the mean square length of chains in unstrained state. *k* is the Boltzmann constant and N refers to the number of chains in the network. This term is much smaller in magnitude as compared to the 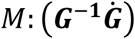 term and is hence neglected in the remaining calculations.

### 5. Dynamics of SF remodeling under cyclic stretch

Uniaxial cyclic stretching of cells causes reorientation along a near-perpendicular direction to the direction of applied stretch. The reorientation dynamics is dependent on the stretch amplitude as well as the duration of stretch. Based on our observations, we propose a reorientation law for the SF (7,12)

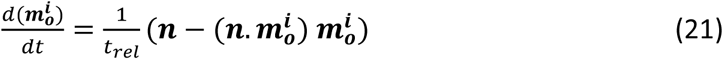

The vector **n** represents the unit vector along the direction of preferred cytoskeletal orientation under stretch, which is along Y-axis (**e**_2_) (when the cyclic stretch is applied along X-axis). The term *t_rel_* refers to a remodeling time scale which also depends on the amplitude of cyclic stretch. Using this reorientation law, we can predict the remodeling dynamics of actin stress fiber in cells under different stretch amplitude and duration. Using the equations for 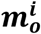 the reorientation law becomes

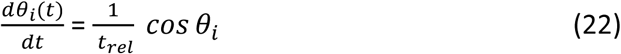

### 6. Changes in the effective Young’s modulus of the cell due to stress fiber remodeling

We quantified the effective Young’s modulus along the direction of SF reorientation (i.e ***e***_2_) as the gradient of uniaxial tension, N, with respect to the stretch in that direction (10)

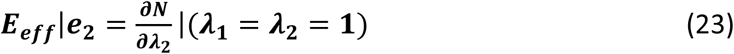

Where N= *t*_2_ *and t*_1_ = *t*_3_ =0. From the constitutive equation, we can therefore write the tractions along the ***e***_1_, ***e***_2_ and ***e***_3_ direction as

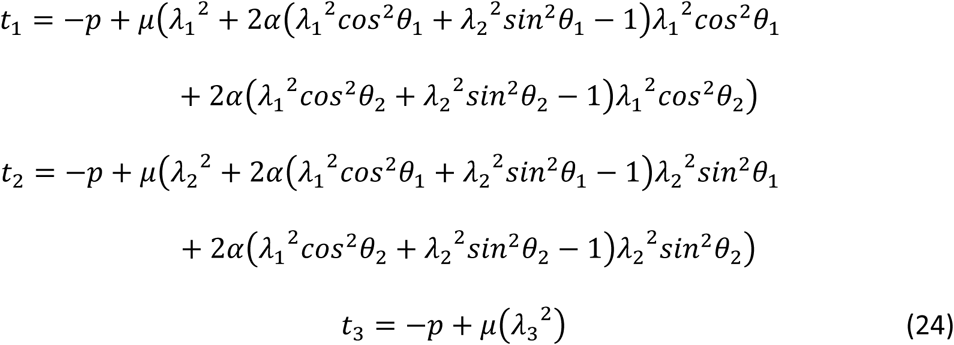

Using the expressions of *t*_1_, *t*_2_ and *t*_3_, we write two additional terms as

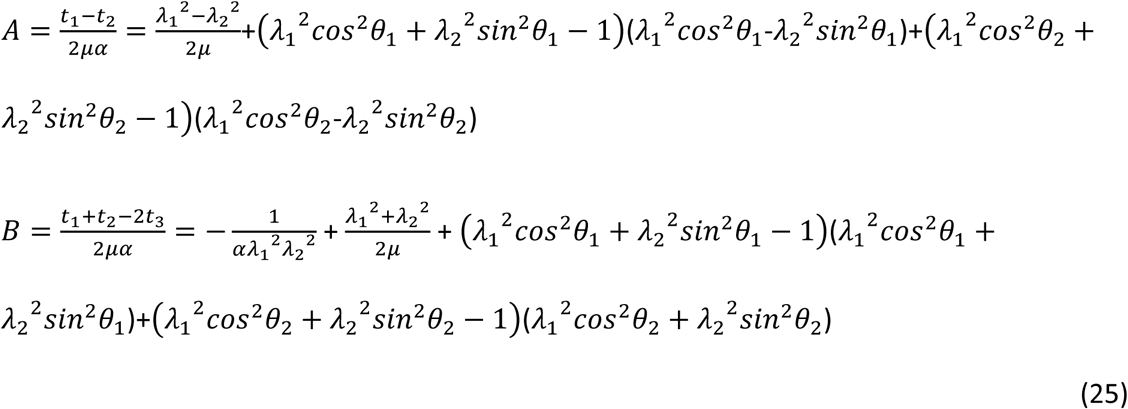

Using the expressions of 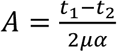 and 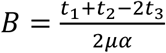 we compute the term 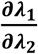 and using equation (23) we obtain the expression of

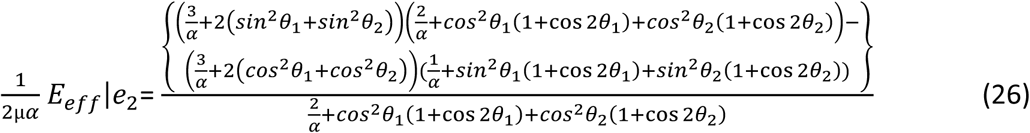

From this expression at *θ*=90°; we obtain *E_eff_*|***e***_2_=*μ*(3 + 8*α*). The parameters µ and α are obtained by equating with the experimentally obtained Young’s modulus values obtained from the AFM indentation experiments.

### 7. Formulation of generalized structure tensor based on distributed fiber orientations

We quantified the angular orientation of the distributed SF in the cell using a two term Gaussian distribution function. The orientation distribution function 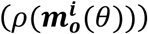 is based on the distribution of fibers in their referential configuration.

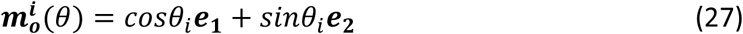

The Gaussian distribution has a symmetry requirement 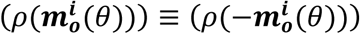 and is also normalized such that

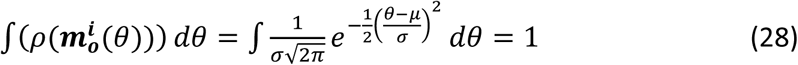

Based on these distributions we formulate a generalized symmetric structure tensor using the relation [13]:

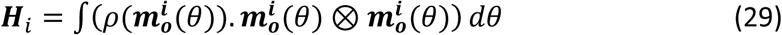

Where *i* ∈ (1,2) for the two families of distributed SF. Components of the structure tensor **H** are calculated as:

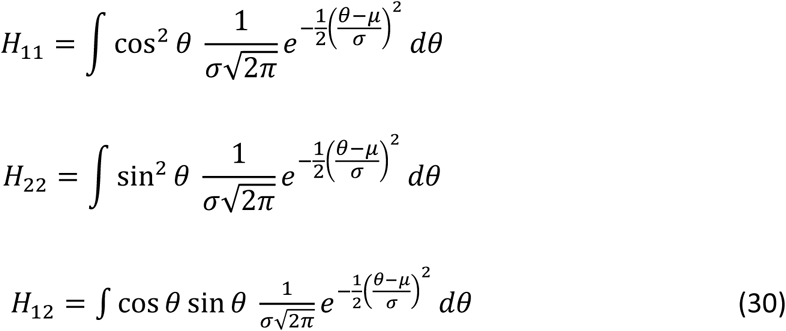

Because **H** is symmetric, *H*_12_ = *H*_21_, and the other terms are zero. Using the structure tensor for the distributed fiber reorientations we write the constitutive equation as:

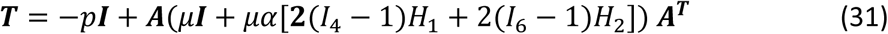

Where *I*_4_ and *I*_6_ can be re-calculated using *H* as

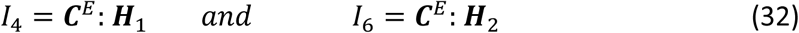

